# Urban living influences the reproductive success of Darwin’s finches in the Galápagos Islands

**DOI:** 10.1101/2020.07.08.193623

**Authors:** Johanna A. Harvey, Kiley Chernicky, Shelby R. Simons, Taylor B. Verrett, Jaime A. Chaves, Sarah A. Knutie

## Abstract

1. Urbanization is expanding worldwide and can have major consequences for organisms, anthropogenic factors can reduce the fitness of animals but may also have benefits, such as consistent human food availability. Understanding these trade-offs is critically important in environments with unreliable annual natural food availability, such as the Galápagos Islands where urbanization is rapidly increasing. For example, during dry climatic condition years, the reproductive success of bird species, such as Darwin’s finches, is low because low precipitation reduces food availability. Urban areas in the Galápagos provide supplemental human food to finches, which could improve their reproductive success during years with low natural food availability. However, urban finches might face trade-offs, as the incorporation of anthropogenic debris (e.g. string, hair, plastic) into their nests can increase mortality.
2. In our study, we determined the effect of urbanization on the reproductive effort and success of small ground finches (*Geospiza fuliginosa*; a species of Darwin’s finch) during a dry year on San Cristóbal Island, Galápagos Islands. We also documented the abundance of anthropogenic debris incorporated in to nests.
3. We quantified nest building, egg laying, hatching, and fledging of small ground finches in an urban and non-urban area. We also qualified the type of anthropogenic debris in finch nests and the quantified the percent of debris comprising total nest mass. We determined whether incorporating these materials into the nest directly led to entanglement- or ingestion-related mortalities.
4. Overall, urban finches built more nests, laid more eggs, and produced more fledglings than non-urban finches. However, every nest in the urban area contained anthropogenic-related material, which resulted in entanglement- or ingestion-related mortalities in 18% of nests with nestlings. Non-urban nests did not contain any anthropogenic-related material.
5. Our study showed that urban living has trade-offs during drier climatic conditions: urban birds have overall higher reproductive success than non-urban birds, but urban birds can also suffer a negative consequence by using anthropogenic-related material for nesting. These results suggest that despite the potential cost of urban living, finches benefit overall from urban living and urbanization may buffer the effects of limited resource availability in the Galápagos Islands.

## 1. Introduction

Urbanization is increasing worldwide with fewer pristine places remaining untouched by humans (Vitousek, 1997). Urbanization can directly change the physical structure of the ecosystem through the creation of roads and buildings and introducing artificial light, pollution, and noise (Fernández-Juricic, 2002; Dominoni, Quetting, & Partecke, 2013; Herrera-Dueñas et al., 2017). Consequently, native fauna can suffer reduced fitness or extirpation in response to urbanization-related stressors, such as environmental change, increased predation, limited natural food availability, and increased disease and parasites (Blair, 1996; Lepczyk, Mertig, & Liu, 2004; Bailly et al., 2016; Johnson & Munshi-South, 2017). However, urban living can also benefit organisms by reducing the natural predation risk and increasing alternative resource availability, such as human food sources and habitat structures (Gering & Blair, 1999; Lowry, Lill, & Wong, 2013; Møller et al., 2015). The effect of anthropogenic materials on seabirds has been well examined (Roman et al., 2019); however, the impacts on passerines, particularly on reproductive success in urban areas, have not been well assessed. The effect of urbanization on birds can vary, but include earlier lay dates and lower reproductive success in urban versus non-urban areas (Chamberlain et al., 2009; Sepp et al., 2018). Urban food availability has been suggested as a principal factor driving the variation of demographic responses across passerines (Chamberlain et al., 2009). While species diversity can decline in urban areas (Kark et al., 2007), urban areas still sustain a number of native species (Aronson et al., 2014); this duality presents an opportunity to better understand the trade-offs experienced by a species in response to urbanization.

Determining the effects of urbanization on islands is especially important given that islands host higher human population densities, as compared to the mainland, and harbor 20% of all terrestrial plant and vertebrate species diversity (Kier et al., 2009; Courchamp et al., 2014). Furthermore, island endemic species, particularly specialist species, can be highly sensitive to natural and anthropogenic perturbations (Buckley & Jetz, 2007) due to their small population sizes, low immigration, and associated genetic factors, such as low genetic diversity (Benning et al., 2002). However, the effect of urbanization on island species can vary. For example, native Caribbean island reptiles can persist in urban environments, but in lower numbers (Jesse et al., 2018). In contrast, urban adapted dark-eyed juncos have longer breeding seasons, which result in higher reproductive success (Yeh & Price, 2004). These few studies suggest that native species can differ in their responses to urbanization. As island species face extinction threats on many fronts, examining trade-offs for urban animals on islands could provide insight into their ability to respond to anthropogenic change or help inform management and conservation of the species.

The Galápagos Islands of Ecuador has experienced recent urbanization due in large part to growth in ecotourism. Since the 1990s, Galápagos tourism has increased by an average of 9.4% per year, with current estimates of nearly 225,000 visiting tourists each year. The resident human population has increased by an average of 6.4% per year since the early 1990’s, reaching 25,244 in 2015 (Epler, 2007; Walsh & Mena, 2016). The recent human population growth and the incipient urbanization of the Galápagos islands provides an ideal “laboratory” to determine the effects of human activity on endemic animals. For example, recent studies have shown that Darwin’s finches in urban areas prefer non-natural food compared to finches in non-urban areas (De León et al., 2018), which consequently resulted in changes to their microbiota (Knutie, Chaves, & Gotanda, 2019), epigenetics (McNew et al., 2017), and morphology (Hendry et al., 2006; De Léon et al., 2011).

The Galápagos also face natural stressors, such as highly variable climatic conditions. The islands have a hot, wetter season from approximately January to May, and a cool, drier season from approximately June to December (P. R. Grant & Boag, 1980). The conditions during these seasons depend on the Inter-Tropical Convergence Zone (ITCZ) and the periodically irregular El Niño Southern Oscillation (ENSO) (Trueman & d’Ozouville, 2010). El Niño events can often result in wetter seasons with high primary productivity and therefore high food resources for the finches, whereas La Niña events are characterized by drier seasons with limited primary productivity and food resources (P. R. Grant & Boag, 1980; Trueman & d’Ozouville, 2010). Consequently, low reproductive fitness has been a consistent documented pattern reported in Darwin’s finches across the Galapagos islands in dry La Niña years, with this effect being more pronounced in the arid coastal zones (Boag & Grant, P.R., 1981; Gibbs & Grant, P.R., 1987; Grant & Grant, 1989, 1999; Koop, LeBohec, & Clayton, 2013). Low reproductive success in response to dry years has also been found in other island land birds, such as Galápagos mockingbirds (Curry & Grant, P.R., 1989; McNew et al., 2019). To date, studies have not examined the influence of urbanization on the reproductive fitness of Darwin’s finches during dry conditions, Therefore, the Galápagos islands present a unique opportunity to examine the effects of growing, yet incipient, urbanization in a landscape where climate could be further exacerbating the positive or negative effects of urbanization on an endemic species.

In our study, we examined the effect of urbanization on the reproductive effort and success of small ground finches (*Geospiza fuliginosa*; a species of Darwin’s finch) during a year with dry climate conditions. First, we determined whether reproductive effort (i.e. nests built, eggs laid, hatchlings) and success (i.e. young fledged) of small ground finches differed between urban and non-urban areas by tracking the survival of nests from construction to egg laying, hatching, nestling survival, and confirmed fledging of young. During years with dry conditions, the reproductive effort and success of Darwin’s finches is lower than years with wet conditions, which has been linked to reduced natural food availability (Boag & Grant, P.R., 1981; Koop, LeBohec, & Clayton, 2013). Because urban areas are supplemented with additional human food resources (De León et al., 2018), finch reproductive effort and success is predicted to increase in urban areas compared to non-urban areas. However, finches incorporate human-related material into their nest (Knutie et al., 2014), which is readily available in urban areas and can result in death due to entanglement (Townsend & Barker, 2014; Jagiello et al., 2018; Jiguet et al., 2019, Theodosopoulos & Gotanda, 2019). Therefore, although urban finches are predicted to have higher overall reproductive success, urban finches likely face a trade-off related to anthropogenic debris in their nests: urban finches are predicted to incorporate more anthropogenic debris into their nests and suffer higher mortality related to this debris (i.e. entanglement, ingestion) than non-urban finches.

## 2. Materials and Methods

### 2.1 Study system

We conducted our study between February – May 2018 (during the breeding season) in the arid lowland climatic zone of San Cristóbal (557 km^2^) in the Galápagos Islands. Breeding for ground finches is initiated by heavy rainfall events and continued breeding is dependent on continued rainfall (Boag & Grant, P.R., 1984; S. Kleindorfer, 2007). Rainfall on San Cristóbal is highly variable, with interannual variation alternating between high and low rainfall (P. R. Grant & Boag, 1980).

We quantified nest building, egg laying, hatching, and fledging of small ground finches in an urban and non-urban area. The urban area was in the capital city of Puerto Baquerizo Moreno (hereon, urban area), which is the second largest city in the Galápagos archipelago with a human population of 7,199 (INEC, 2016). The urban area primarily consists of impermeable concrete or stone surfaces and human built structures in altered landscapes. Our urban study area measured 0.79 km^2^ (∼1.2 km by 0.62 km) and included tourist and residential zones (Fig. 1C). The search area within the urban area was determined due to suitability of habitat, i.e. presence of trees for nesting and accessibility, permission to search and access granted to private properties. The non-urban area was in the Jardin de Opuntias (hereon, non-urban area), which is a Galápagos National Park site located 4.5 km southeast of the urban area consisting of vegetated natural habitats with no unnatural impermeable surfaces present. Our non-urban study area measured 0.21 km^2^ and covered 1.4 km of the main trail and 0.15 km to each side (Fig. 1D). The search area is larger in the urban area than the non-urban area due to spatial mismatch and differences in environmental structure, which can result in urban patches devoid of suitable nesting areas. Search efforts, via person-hours (total number of hours searched per person for each day), were tracked for each study area to normalize search efforts. Our search area was primarily, but not exclusively centered, by the presence of the arboreal cactus, *Opuntia megasperma*, which are one of the preferred nesting locations of small ground finches. Cacti are rare on San Cristóbal, likely due to destruction by introduced mammals in the 1800s, but are found in high abundance within the Jardin de Opuntias (Phillips, Wiedenfeld, & Snell, 2012; Dvorak et al., 2019). Small ground finch nests are commonly found in cacti as well as trees such as matazarno (*Piscidia cathagenensi*) and Galápagos acacia (*Acacia rorudiana*). The non-urban area receives very low human visitation: locals occasionally, but rarely, visit the site to access the beach. Cacti are also frequently cultivated in urban areas, and therefore are found in garden beds, planters, city parks, and the main boardwalk. Urban finches also nest in native and non-native trees and occasionally nest in human built structures, such as gutters and building signs.

**Fig. 1.**
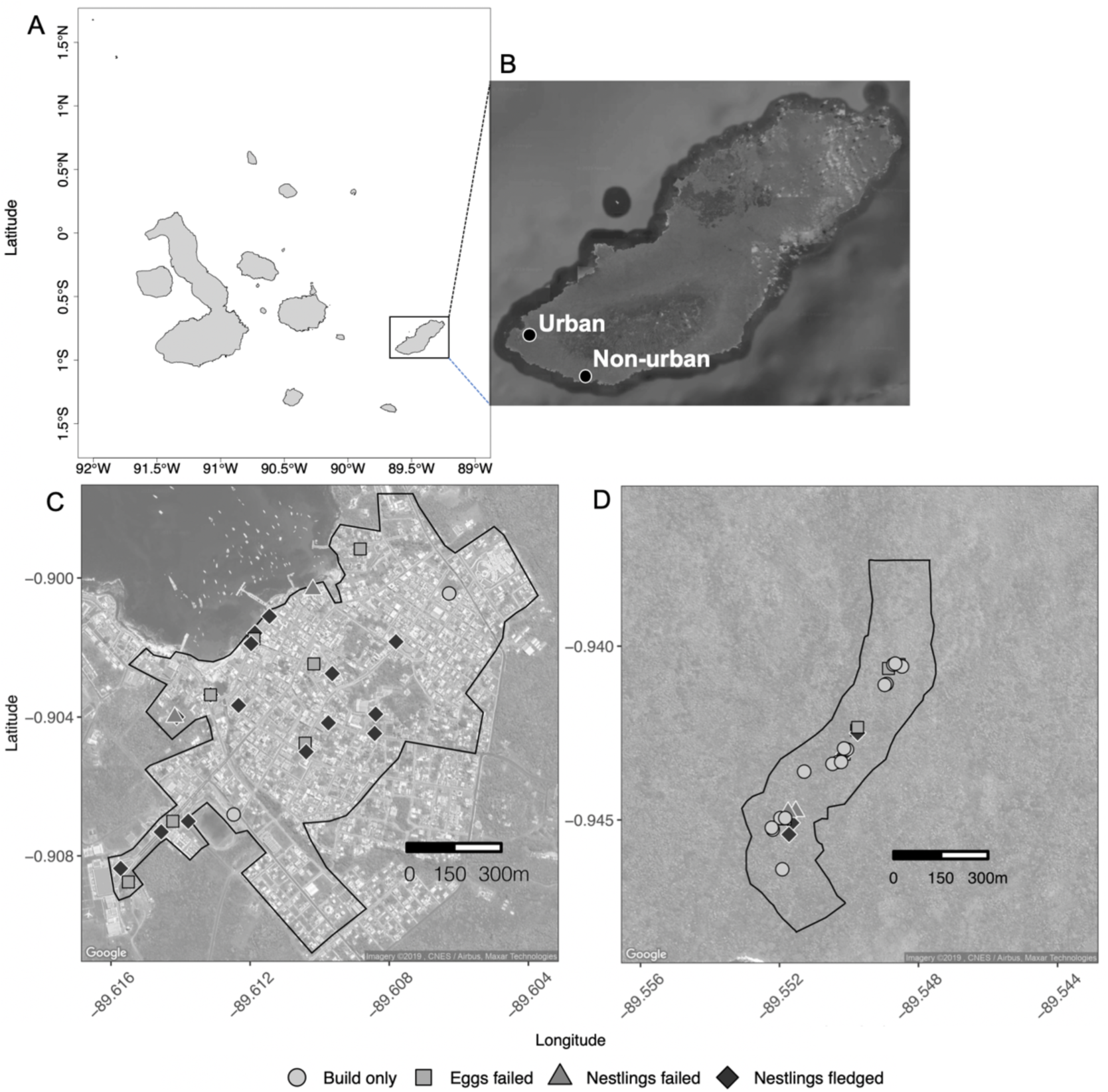
(A) Map of the major islands of the Galápagos archipelago and inset map of (B) San Cristóbal Island with sampling areas noted (black dots) for the urban area (Puerto Baqueizo Moreno) and the non-urban area (Jardin de Opuntias). Satellite maps of the (C) urban area and (D) the non-urban area showing nests which were builds only (grey circle), nests with eggs that failed (grey square), nests with nestlings that failed (dark grey triangle), and nests with nestlings that fledged (black circle) across each sampling site with the search area delineated by the black polygon border. Map data Google Maps Imagery © 2019 and the GADM database (Hijmans et al., 2014).

### 2.2 Locating nest sites and data collection

In each urban and non-urban area, we searched intensively for nests and for small ground finches exhibiting nest-building behaviors, including vocalization and behavioral cues. The field sites were searched nearly every other day for evidence of nest-building activity by small ground finches. We followed all nests accessible with the use of a 10-foot (∼3 meter) ladder. Once found, nests were checked every other day until eggs were found in the nest. Observations were made primarily through binoculars to minimize nest disturbance, and secondarily through a small camera (Contour LLC, Provo, USA) attached to an extendable pole when the nest was not attended by adults. Once the eggs hatched, we followed the survival of nestlings and banded them with a unique color band combination when they were 7-8 days of age (hatch date = day 0). Successful fledging was confirmed by identifying individual birds once they left the nest, as in previous studies (Knutie et al., 2016). After nestling birds fledged or died, the nest was collected and placed in a sealed plastic bag. Each nest was carefully dissected to separate natural and anthropogenic materials, after which each material type was weighed (g). Anthropogenic materials were then qualitatively identified (composition of material, possible source, and color of materials) when possible. All detected nest failures and mortalities were documented, and causation was determined when possible. Materials associated with mortality via ingestion or entanglement were also identified and documented.

Our study resulted from a single year of sampling, and no building or breeding individuals in our study had been previously banded. We did not observe any dispersal of banded birds across urban and non-urban areas. The flight distance between urban and non-urban sampling areas is eight km, and previous studies have not found dispersal in small ground finches to occur across habitats (Sonia Kleindorfer et al., 2006). Therefore, it is unlikely that small ground finches disperse between the urban and non-urban area of this study at significant rates.

### 2.3 Statistical analyses

All data were analyzed in Rstudio v1.2 (R Core Team, 2012). We calculated total person-hours by multiplying the number of hours searched by the number of people searching for each day at both sites (urban and non-urban respectively) across all days of the survey period. We tested survey effort using an independent t-test on total person-hours per day at each site after examining data for homogeneity of variance using a Fligner-Killeen test and Q-Q plots for assessment of normality. For effect size, we calculated Cohen’s D (Cohen, 2013) for percent of nestlings fledged per nest across sampling areas and percent nest mass of anthropogenic debris across sampling areas, respectively. Cohen’s D is a standardized measure of difference in means divided by the pooled standard deviation, and it was calculated using the lsr package (Navarro & Foxcroft,).

We used generalized linear models (GLMs) using the glm function in the lme4 package (Bates et al., 2015) and ANOVAs using the car package (Fox & Weisberg, 2011). We first verified that data met assumptions of models by checking for overdispersion and underdispersion. We used GLMs with a binomial error structure to determine whether location (urban and non-urban) affected the binary presence or absence of nests with eggs, nests with nestlings, and trash in nest material. Additionally, we used binomial logistic regressions, which are a special case of GLM to determine the effect of location on hatching and nestling survival trials, independently. The response variable is a matrix of trials, which are successes and failures, where the number of eggs hatched and nestling fledged per nest represent successes while eggs not hatched and nestling mortalities represent failures, respectively. We also used a binomial logistic regression to determine the effect of location along with the proportion of anthropogenic debris in the nest on nestling survival, where nest survival was measured as successes and failures.

We used a Mayfield logistic regression, an extension of the Mayfield method, which examines nest survival data over the incubation and nestling periods and incorporates explanatory variables in a logistic regression (Hazler, 2004). Nest fates (success or failure) are modeled over the number of exposure days where each exposure day is a trial. The explanatory variables are covariates of location (urban and non-urban) and proportion of anthropogenic debris in nests which are also weighted by the number of exposure days (Hazler, 2004). Exposure days were calculated using the median of the last day the nest was known active to the last check day when the nest outcome or fate was known (Mayfield, 1961).

## 3. Results

Nest searching was conducted over 49 days in the urban area with an average of 12.51 ± 4.21 person-hours search day. Nest searching was conducted over 40 days in the non-urban area with an average of 12.23 ± 7.5 person-hours per search day. The number of person-hours spent searching did not significantly vary across the urban and non-urban study areas (Independent t-test, t = -0.22, *df* = 75.51, *P* = 0.82).

The first urban finch nest with eggs was found on February 10, 2018, whereas the first non-urban finch nest with eggs was found on March 7, 2018 (Fig. 2). The last urban nest with eggs was found on April 14, 2018 and the last non-urban finch nest with eggs was found on April 13, 2018, resulting in a breeding season of 68 days in the urban area and 37 days in the non-urban area. Small ground finches built 29 nests in the urban area and 29 nests in the non-urban area (Fig. 1, Table 1). The urban area had more nests with eggs (n = 25 nests with eggs out of 29 nests built) than in the non-urban area (n = 12 nests with eggs out of 29 nests built) (GLM, *χ*^*2*^ = 13.28, *df* = 1, *P* < 0.0001).

**Table 1.** Reproductive success of small ground finches in non-urban and urban areas.

| Variable | Non-urban | Urban |
| --- | --- | --- |
| No. built | 29 | 29 |
| No. with eggs | 12/29 (41.3%) | 25/29 (86.2%) |
| No. with nestlings | 6/12 (50.0%) | 17/25(68%) |
| No. with at least one fledgling | 2/6 (33.3%) | 15/17 (88.2%) |
| No. with anthropogenic materials | 0/10 (0%) | 22/22 (100%) |

**Fig. 2.**
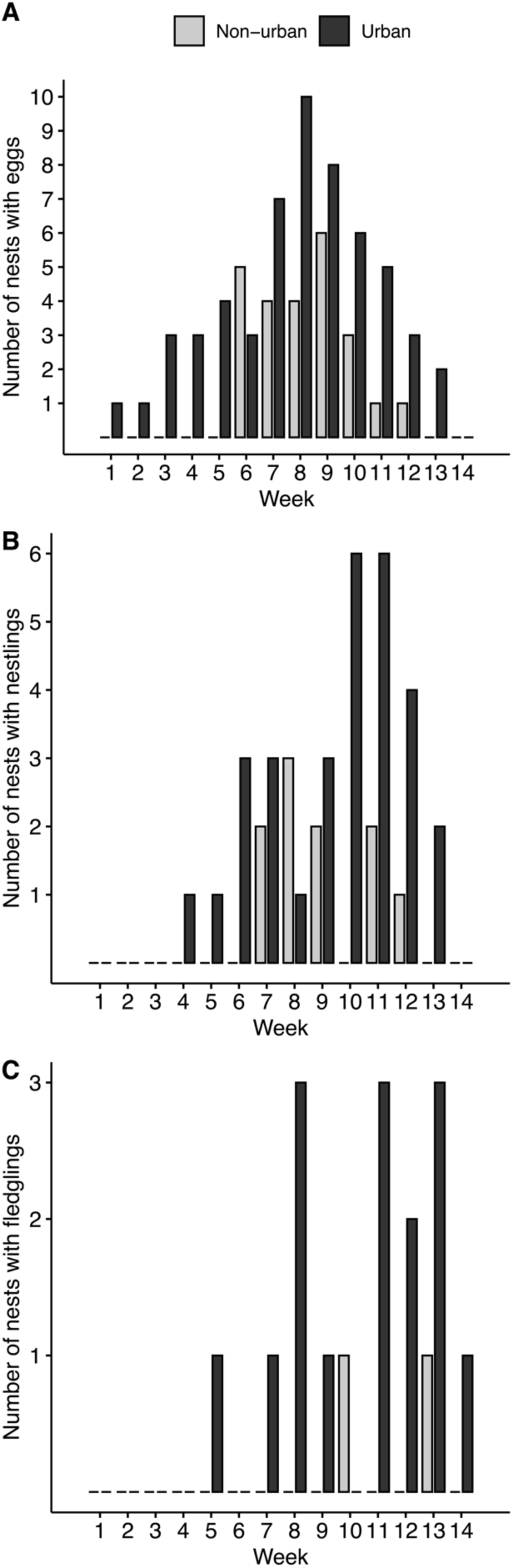
Histograms of reproductive activity noting frequency of nests with (A) eggs, (B) nestlings, and (C) fledgling (where fledging occurred that week) across each week of the study period in non-urban (grey) and urban (black) sampling areas.

The urban area had more nests with hatchlings (n = 16) from the total nests built (n = 29) compared to the non-urban area (n = six nests with hatchlings from 29 nests built) (GLM, *χ*^*2*^ = 9.0, *df* = 1, *P* = 0.003). Of the 25 urban nests with eggs, eight failed at the egg stage. Three nests were damaged during likely predation events, with no remains found and the nest entrance destroyed. Two nests were found abandoned with eggs intact and cold. One nest had only one egg and was found predated by ants. One nest was found with a single egg ejected and the remaining eggs found cold in the unattended nest. One nest was found empty with no evidence of predation or cause of failure determined. Of the 12 non-urban nests with eggs, six failed at the egg stage. One nest had clear signs of predation with a broken egg shell found outside of the nest. One egg nest was found empty with no evidence to explain its failure. The remaining four failed egg nests were abandoned, with cold eggs found in the unattended nest.

Overall survival, from egg stage and fledging, was higher in the urban compared to the non-urban area: urban nests had a higher proportion of nests with eggs that hatched (*χ*^*2*^ = 4.34, *df* = 1, *P* = 0.04) and nestlings that fledged (*χ*^*2*^ = 14.35, *df* = 1, *P* = 0.0002, *Cohen’s D* = 1.48) than the non-urban nests. Only two urban nest failures occurred during the nestling stage. One nest was found with a six-day old dead nestling hanging from ingested hair that was woven into the nest (Fig. 3A) and the two other nestlings missing from the nest. One nest was found empty with no apparent cause for mortality or nestling remains found. For three out of the four nest failures during the nestling stage in the non-urban area, nestlings were missing with no clear signs of depredation. These mortalities occurred post-hatching at days six, seven, and ten days old. One non-urban nest had signs of depredation, with partial nestling remains (i.e. limbs and skull) found near the nest.

**Fig. 3.**
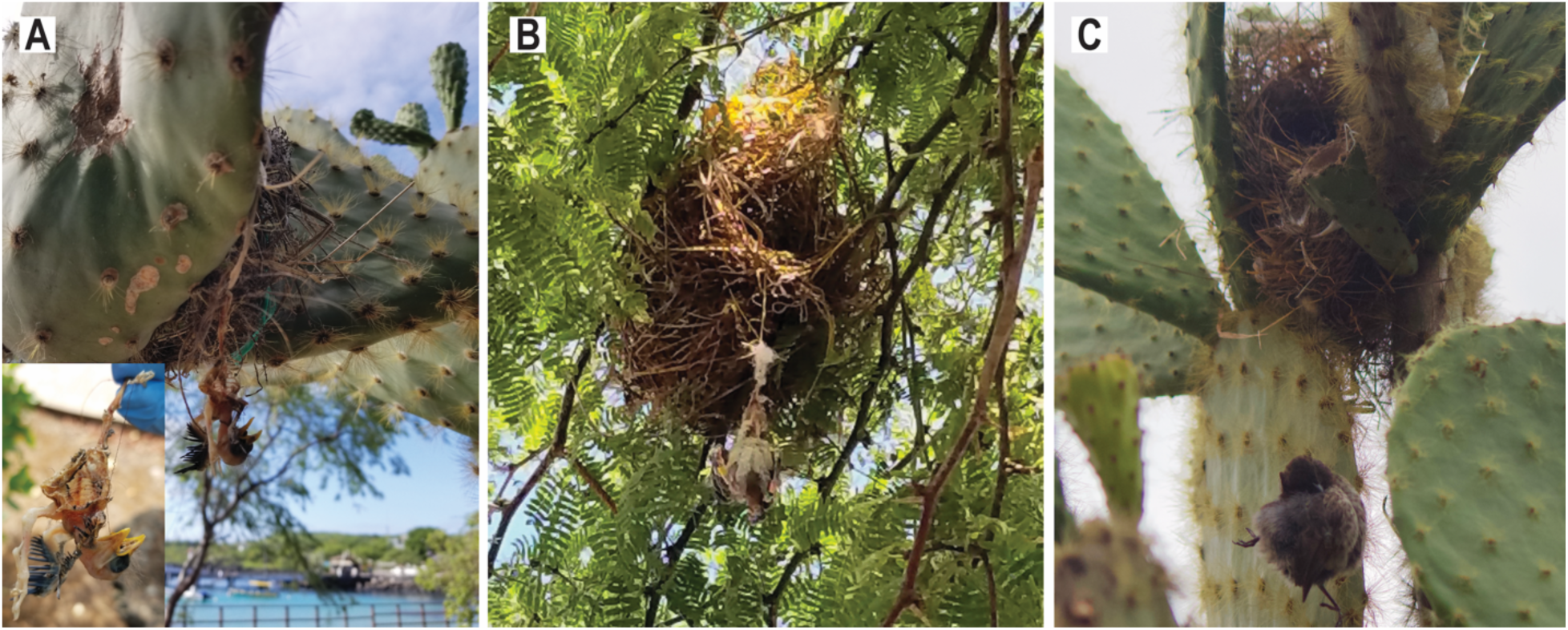
Documented small ground finch mortalities due to ingestion/entanglement of anthropogenic nest materials: a. 6-day old nestling ingestion/entanglement with plastic and human hair, b. 12-day old nestling entanglement with synthetic string, c. adult female entanglement during nest building with human hair).

Anthropogenic debris was not found in non-urban areas, whereas all nests in the urban area contained anthropogenic debris (*Cohen’s D* = 2.140, Table 1, Fig. 4). The percent of anthropogenic debris out of total nest mass varied from 3.1% to 22.7% (Supporting Information). Within the urban area fledging survival, examining the number of nestlings fledged versus nestlings failed, declined with increasing proportion of anthropogenic material comprising total nest mass (GLM, *χ*^*2*^ = 13.80, *df* = 1, *P* = 0.0002). When we jointly model the covariates of location, urban and non-urban, with the proportion of nest trash to determine the effect on daily nestling survival, using a Mayfield logistic regression, we found both covariates to be significant (*β*_*1*_= -1.169, *P* <0.0001, *β*_*2*_= -4.109, *P* = 0.0391).

**Fig. 4.**
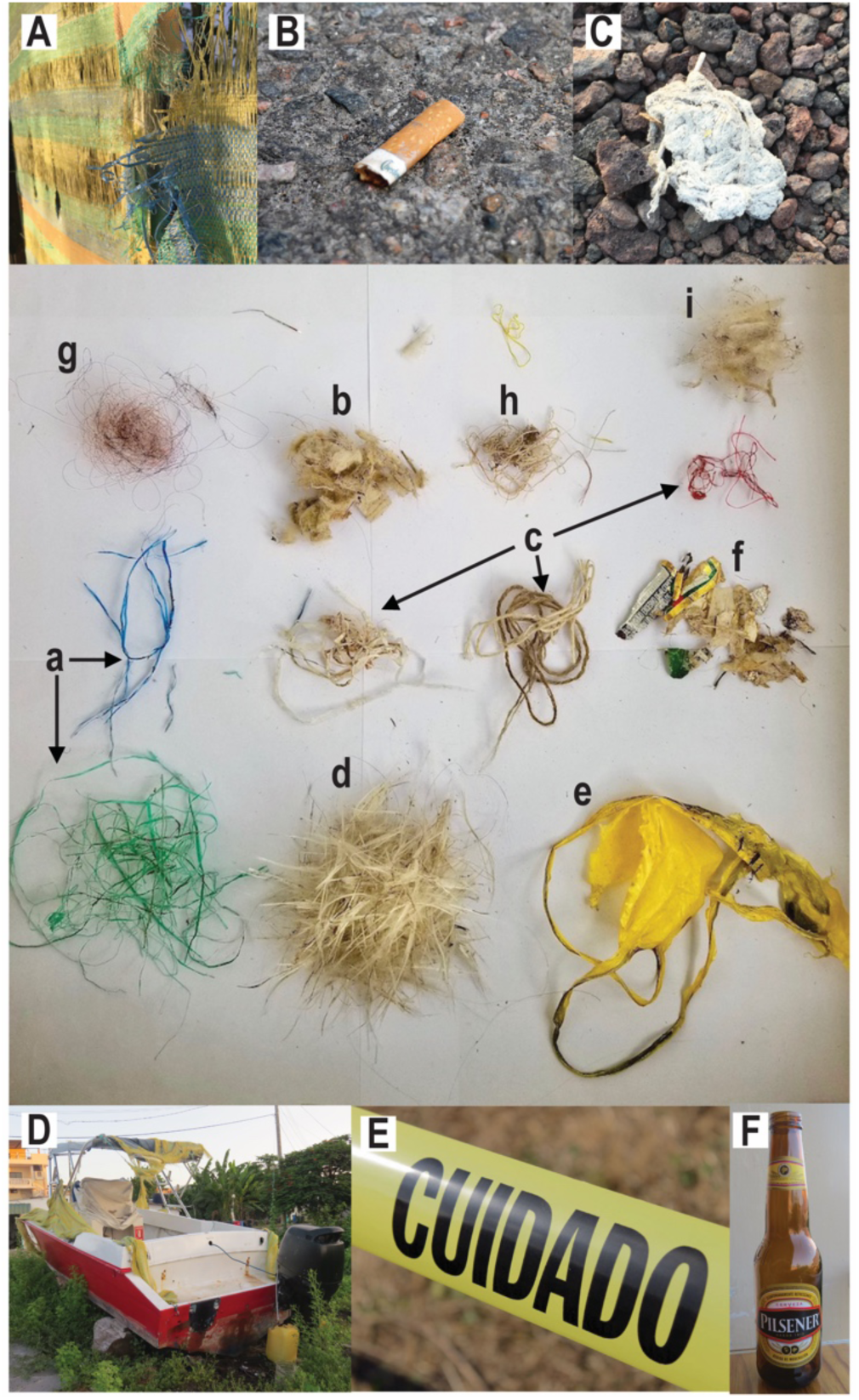
Identification of anthropogenic materials dissected (a. shredded plastic tarp strands, b. cigarette fibers, c. fibers/thread, d. fiberglass from fishing boats, e. caution tape, f. paper shreds, g. human hair, h. shredded twine, i. synthetic stuffing) from a single *G. fuliginosa* urban nest which were sorted by material type and color (central figure, labeled with lower case) with potential urban source materials shown (below and above, labeled with upper case).

The most common materials found across nests were synthetic strings and fibers and synthetic stuffing followed by plastic (Supporting Information, Fig. 4). The color of anthropogenic nest debris most collected was white (present in all nests), followed by blue/purple and green. In the urban area, we found and documented four cases of nest entanglement-associated mortalities, which include three nestlings from separate nests and one adult female, directly related to anthropogenic debris: 1) one nestling was found hanging from ingested hair that was woven into the nest, with its leg also tangled in free hanging plastic string (Fig. 3A), 2) one nestling was hanging from ingested synthetic string (Fig. 3B), 3) one nestling (near fledging) was found with its leg entangled in plastic string, and 4) one adult female was found hanging, strangulated, from human hair during active nest building (Fig. 3C).

## 4. Discussion

Low reproductive success in dry years is an established pattern in Darwin’s finches. The effect of dry years on the reproductive success of urban Darwin’s finches, however, has not been previously addressed. In this study, we found, that consistent with the majority of urban rural comparisons (Chamberlain et al., 2009) urban small ground finches’ reproductive effort, egg laying, began earlier then in their rural congeners. However, reproductive effort as well as reproductive success was significantly higher, in urban finches compared to non-urban finches: the urban area resulted in higher numbers of nests with eggs (Fig. 2A), nests with nestlings (Fig. 2B), and fledging success (Fig. 2C). All urban finches were found to incorporate anthropogenic debris (3.1% to 22.7% of total nest mass) into their nests while non-urban nests had no anthropogenic debris incorporated. Critically, mortalities due to anthropogenic nest debris entanglement were recorded across four urban nests, affecting 18% of nests with nestlings and one female in active nest building, and shown to be a cost associated with nesting in urban areas. These results suggest that despite anthropogenic debris related mortalities, small ground finches derived an overall reproductive benefit from urban habituation during a dry year, perhaps associated with higher reproductive output due to earlier initiation of the breeding season and higher offspring survival, which may be attributable to the high urban food availability.

The later initiation of breeding and reproductive failures documented in non-urban finches are likely associated with limited precipitation and dry La Niña conditions as seen previously (Boag & Grant, 1981, 1984; Koop, LeBohec, & Clayton, 2013). Rainfall and the associated increase in primary productivity is known to initiate and sustain finch breeding (Gibbs & Grant, P.R., 1987; Zhang et al., 2019; MEI.v2 Retreived 21 December 2019)(Boag & Grant, P.R., 1984). Indicating enough rain occurred to trigger breeding but perhaps not enough natural food resources were available to sustain breeding efforts. During the 2018 breeding season, we recorded a high percent of nest abandonments in the non-urban area: 33% of finch nests (four of 12 nests) were abandoned and failed with eggs. The lower limit of rainfall required to sustain finch breeding is unknown and may also be impacted by rainfall the previous year and the associated carryover effects on vegetation and invertebrate communities. The previous year, 2017, was a milder La Niña year (Zhang et al., 2019), providing some support for the premise of carryover of low primary productivity further affecting the availability of resources in 2018. The differences in overall reproductive output of urban and non-urban small ground finches in our 2018 study might be related to the higher food resource availability in the urban area (Lochmiller & Deerenberg, 2000), and low primary and secondary production associated with low precipitation typical of a La Niña year in the non-urban area. Urban food resources are independent of climatic variability, whereas natural food resources (seeds and insects) are both lower in dry years (low precipitation, drought). Accordingly, Galápagos land nesting birds have shown reduced breeding success in dry years (Grant & Grant, 1993; Koop, LeBohec, & Clayton, 2013; McNew et al., 2019). Non-urban finch fitness, in terms of both breeding success and adult survival, is dictated by precipitation patterns and the resulting food availability (Gibbs & Grant, P.R., 1987; Koop, LeBohec, & Clayton, 2013; Grant & Grant, 2014). Therefore, future studies should examine whether the urban breeding success exceeds non-urban success in wet years and across years in order to understand the long-term demographic patterns of urban finches. Urban and non-urban birds seem to face distinct and at least partially non-overlapping stressors; non-urban birds are more impacted by climatic variables, and resulting food availability, and urban birds are being impacted by anthropogenic stressors, such as anthropogenic debris and possibly low nutritional food quality.

While human-based food availability in urban environments can benefit adult birds, nestlings require a higher protein diet (Boag, 1987). Natural high-protein food sources, such as arthropods, demonstrate declines with increasing urbanization (Shochat et al., 2004). Additionally, the lower quality of food in urban environments has been seen to negatively affect nestling and juvenile growth in other urban bird species (Pierotti & Annett, 2001; Seress et al., 2012, 2020). Even if urban food quality is lower nutritionally than natural food sources, the quantity and consistent availability of the resources can provide short-term benefits for survival of urban finch nestlings. However, poor nestling diet can directly impact morphological metrics later in life (Boag, 1987), indicating possible lasting changes in urban finch adults. Urban finches were observed foraging on a variety of human foods (i.e. chips, rice, bread, corn), but also seen foraging on refuse (i.e. out of garbage bins, rotting organic material in street) (pers. obs. JAH, TBV). While previous studies have examined the diet of finches in urban vs. non-urban areas (De León et al., 2018), an examination is still needed of the resulting effect of low quality diet on urban finch on long term trends in demography. Furthermore, future studies are needed to determine the long-term effects of urban nestling diet on adult morphology and overall fitness. The marked variation between urban and non-urban breeding success found here may have also been compounded by climatic factors which impact resource availability and these differences, such as parental provisioning rates and diet analysis, must be taken into account in future studies.

We found that 100% of urban nests contained anthropogenic debris, whereas no anthropogenic debris was recovered from non-urban nests. These results were likely due to debris availability: abundant anthropogenic debris was available in the urban area and only two pieces of debris were seen at the non-urban area during the field season. Furthermore, 24% of urban nests were associated with debris-related mortalities and the proportion of anthropogenic debris comprising the total nest mass impacted nestling survival. Other studies have observed mortalities due to both entanglement and ingestion of anthropogenic debris in nestling and adult land birds (Mee et al., 2007; Henry, Wey, & Balança, 2011; Theodosopoulos & Gotanda, 2019). However, few studies have examined the effect of debris on reproductive success in land birds (Jagiello et al., 2018) and only one previous study has examined passerines (Townsend & Barker, 2014). Several types of common anthropogenic debris (e.g., plastic string, human hair, and string fibers) are frequently used by urban finches in nest building and pose a higher risk, as seen in the documented entanglement mortalities. The incorporation of anthropogenic debris into nests presents a clear cost to birds nesting in urban habitats; however, despite suffering mortalities due to entanglement, urban nestlings demonstrated higher survival than non-urban nestlings.

Our study was conducted in an area with intermediate, albeit increasing, urbanization, relative to the other human-inhabited Galápagos Islands. We do not know if a higher degree of urbanization would yield the same beneficial result for finches. A number of species succeed and are abundant at intermediate levels of urbanization and disturbance (Ausprey & Rodewald, 2011; Stracey & Robinson, 2012; Perrier et al., 2018). However, species could reach thresholds at which they no longer benefit from increased urbanization because of further changes in habitat structure which result in a loss of resources (water, food resources, perches, nesting sites) (Blair, 1996; Lee et al., 2004); a threshold of urbanization that is detrimental for Darwin’s finches might not have been reached yet. In addition to direct anthropogenic threats, Darwin’s finches in both urban and non-urban areas are also affected by an introduced nest parasite, *Philornis downsi*, which can affect their reproductive success (Knutie et al., 2016; Sonia Kleindorfer & Dudaniec, 2016; McNew & Clayton, 2018). We did not consider possible effects of *P. downsi* in our study because we lacked the power to detect an effect of the parasite (due to the low reproductive success in the non-urban area). Future studies could examine whether the costs and benefits of urbanization for the finches vary across islands with different levels of human population sizes, as well as study the interactions between urbanization and *P. downsi* in limiting reproductive success of Darwin’s finches.

## Acknowledgements

We thank Karla Vasco for her assistance and logistical support. We would like to thank the Galápagos Science Center and the Galápagos National Park for support. We also thank Kiyoko Gotanda, Sabrina McNew and members of the Knutie lab (Alyssa Addesso, Lauren Albert, Anna Sjodin, Grace Vaziri, and Mackenzie Watkins) for comments on the manuscript. Galápagos photos credited to SAK, JAH, SRS, and TBV. Additional photos credited to Monica Hinkle and Santeri Viinamäk.

## Supporting Information

Table of qualitative data for nest material dissections is available as supplemental document.

## Author contributions

JAH and SAK conceived the study, conducted the data analyses, and wrote the manuscript; JAH, SAK, KC, SRS, TBV collected data; JAC provided logistical support. All authors revised and approved the manuscript.

## Data availability statement

All raw data are available on Figshare (doi provided upon acceptance).

## Ethical statement

All applicable international, national, and/or institutional guidelines for the care and use of animals were followed. All bird handling and work was conducted according to approved University of Connecticut IACUC (Institutional Animal Care and Use Committee) protocols (No. A17-044). Our work was done under GNP permits PC 03-18 and Genetic Access permit MAE-DNB-CM-2016-0041.

